# Spontaneous Facet Joint Osteoarthritis in NFAT1-Mutant Mice

**DOI:** 10.1101/2021.02.23.432595

**Authors:** Jinxi Wang, Qinghua Lu, Matthew J. Mackay, Xiangliang Liu, Yi Feng, Douglas C. Burton, Marc A. Asher

**Author notes:** **Correspondence to:** Jinxi Wang, MD, PhD, Department of Orthopedic Surgery, University of Kansas Medical Center, 3901 Rainbow Boulevard, MS #3017, Kansas City, KS 66160, USA. This paper is dedicated to the memory of Marc A. Asher, M.D. who has passed away after his lifelong devotion to orthopedic surgery and research.

## Abstract

**Objectives:** Although rodent models of traumatically or chemically induced intervertebral facet joint osteoarthritis (FJOA) were previously described, the characteristics of spontaneous FJOA animal models have not been documented. This study aimed to identify the characteristics of a murine model of spontaneous FJOA and its underlying mechanisms.

**Methods:** The lumbar facet joints of mutant mice carrying a disrupted NFAT1 (nuclear factor of activated T cells 1) allele and of wild-type control mice were examined by histochemistry, quantitative gene expression analysis, immunohistochemistry, and histomorphometry using a novel FJOA scoring system at 2, 6, 12, and 18 months of age. The reproducibility of the FJOA scoring system was analyzed by inter-observer and intra-observer variability tests. Tissue-specific histomorphometric and gene expression changes were statistically analyzed.

**Results:** NFAT1-mutant facet joints displayed dysfunction of articular chondrocytes and synovial cells with aberrant gene and protein expression in cartilage and synovium as early as 2 months, followed by osteoarthritic structural changes such as articular surface fissuring and chondro-osteophyte formation at 6 months. Deeper cartilage lesions, synovitis, separation of cartilage from thickened subchondral bone, and tissue-specific molecular and cellular alterations in NFAT1-mutant facet joints became evident at 12 and 18 months. Osteoarthritic structural changes were not detected in wild-type facet joints at any ages, though age-related cartilage degeneration was observed at 18 months.

**Conclusions:** Using NFAT1-mutant mice, this study has identified for the first time an animal model of spontaneous FJOA with age-dependent osteoarthritic characteristics, developed the first FJOA scoring system, and elucidated the molecular mechanisms of NFAT1 mutation-mediated FJOA.

## INTRODUCTION

The intervertebral facet joints (zygapophyseal joints) are in the posterolateral aspect of the vertebral column at each vertebral level (except C1-C2) and play an important role in stabilization of the spine. The articulation of the facet joint is formed between a medially facing superior articular process of the lower vertebra and a laterally facing inferior articular process of the upper vertebra. Unlike the intervertebral disc joints which do not consist of synovial tissue, the paired facet joints are the only true synovial joints in the spine. Almost all the facet joints contain meniscoid synovial folds consisting of the synovial lining cells (synovial intima) and the subintimal stroma, forming a crescent-shaped projection (insertion) into the joint space. The subintimal stroma that lies underneath the synovial intima is composed of loose connective tissue containing primarily fibroblasts, macrophages, blood vessels, lymphatics, and nerves.^1–6^ The facet articular surfaces show varying degrees of inclination in different spinal regions, which determine the directions and ranges of segmental motion. The facet joint capsule is inserted into the margins of the articular facets and is often firmly attached to the ligamentum flavum, thereby restricting rotation and dorsal displacement during extension. Based on these structural and functional characteristics, facet joint disorders including facet joint osteoarthritis (FJOA) are thought to be a common cause of back and neck pain. ^5–8^

FJOA is widely prevalent in middle-aged and older populations, although it is far less studied than other osteoarthritis (OA) phenotypes (e.g. knee and hip OA) and intervertebral disc degeneration (IDD). The overall prevalence of FJOA increases with age, suggesting that age is a major risk factor in its development.^7–10^ Other risk factors, such as sex, obesity, IDD, spinal injury, paraspinal muscle degeneration, occupational factors, smoking, and overexpression of proinflammatory cytokines and specific microRNAs are also proposed to be associated with FJOA. However, most risk factors for human FJOA have been proposed through cross-sectional association studies rather than mechanistic investigations with definitive conclusions. The root cause or precise etiopathogenesis of FJOA remains unclear.^7,11–14^

To understand the pathogenesis of FJOA, several rodent models of FJOA were generated by invasive local manipulations such as intra-articular injection of chemical agents including collagenase, plasminogen activator, and monosodium iodoacetate (MIA) ^15–17^, or through direct traumatic/mechanical injury to the articular surface or joint structure.^18–20^ Those chemically or mechanically induced animal models have improved our understanding of morphological and metabolic changes in post-traumatic FJOA. However, the etiology and unilateral characteristics of those invasive FJOA animal models with acute inflammation and rapid disease progression are distinct from that of spontaneous FJOA in humans that usually occurs bilaterally with slow disease progression. There is little transcriptional (gene expression) similarity between chemically-induced OA cartilage and spontaneous OA cartilage.^21^ To explore the pathogenesis and therapeutic strategies for human FJOA, it is critical to identify and characterize an animal model of spontaneous (non-chemical and non-traumatic) FJOA mimicking human spontaneous FJOA.

NFAT1 (NFATc2/NFATp) is a member of the nuclear factor of activated T cells (NFAT/Nfat) family of transcription factors. Previous studies showed that mice with a disruptive mutation in the NFAT1 binding domain (NFAT1-mutant mice) displayed dysfunction of immune cells and articular chondrocytes with premature OA-like changes in the hip and knee joints.^22–25^ However, the effect of the NFAT1 mutation on synovial joints of the spine was previously unknown. Here we report, for the first time, the identification of age-dependent histologic and molecular characteristics of spontaneous FJOA in NFAT1-mutant mice as well as the development of the first FJOA scoring system.

## RESULTS

### Spontaneous, slow-progressing FJOA in NFAT1-mutant mice

#### Articular cartilage

At 2 months of age, articular cartilage and other joint structures were microscopically visible in both wild-type (WT) and NFAT1-mutant facet joints though the joint size and synovial cavity appeared to be smaller than that of adult mice, suggesting that the NFAT1 mutation does not affect facet joint development in mice. Histopathology of the lumbar spine samples demonstrated reduced intensity of safranin-O staining (red staining for proteoglycans) in the lumbar facet joint cartilage of the NFAT1-mutant (*Nfat1*^−/−^) mice compared to that of the WT mice (***Figure 1**, 2-month*). OA structural changes were not evident at this stage. At 6 months, mild to moderate OA structural changes (e.g. articular surface fibrillation/clefts, focal loss of cartilage, and chondro-osteophyte formation) occurred in the *Nfat1*^−/−^ lumbar facet joints, but not in WT facet joints. Osteoarthritic cartilage lesions were usually more severe in the *Nfat1*^−/−^ superior facet than the inferior facet in the same joint (***Figure 1**, 6-month*). At 12 months, severe osteoarthritic cartilage changes were observed in both superior and inferior facets in *Nfat1*^−/−^ mice, but not in WT mice (***Figure 1**, 12-month*). At 18 months, *Nfat1*^−/−^ facet joints displayed more severe articular cartilage lesions and structural destructions. In contrast, WT facet joints showed only age-related cartilage degeneration characterized by focal loss of chondrocytes and safranin-O staining with mild articular surface abrasion (***Figure 1**, 18-month*).

**Figure 1.**
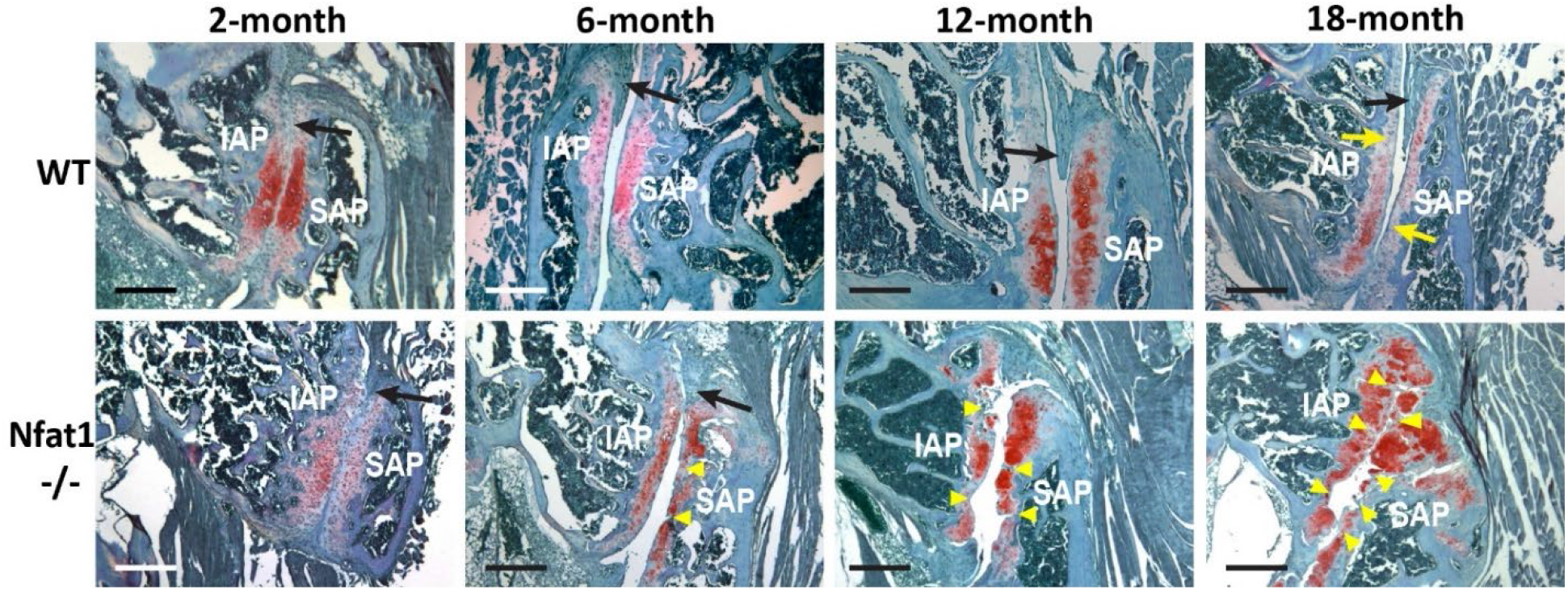
Representative photomicrographs showing age-dependent, slow-progressing osteoarthritic cartilage destructions in NFAT1-mutant (Nfat1−/−) facet joints. Age-matched WT facet joints are presented as controls. Black arrows denote meniscoid synovial folds into the joint space, which are evident in WT joints and Nfat1−/− joints with early OA, but not in the joints with severe OA. Yellow arrows indicate age-related focal loss of Safranin-O staining with mild cartilage degeneration. Yellow arrowheads point to the areas of cartilage lesion. IAP = inferior articular process, SAP = superior articular process. Safranin-O and fast green staining, scale bar = 100 μm.

#### Periarticular tissues

Synovial hyperplasia, characterized by thickening of the synovial tissue due to mild proliferation of intimal synovial lining cells and subintimal connective tissue cells ^26,27^, was observed in some of the *Nfat1*^−/−^ facet joints at 2 and 6 months followed by moderate or severe synovitis in most animals at 12 and 18 months, but not in the WT facet joints at any ages. At 12 and 18 months, ectopic chondrogenic differentiation of the subintimal connective tissue cells with endochondral ossification was observed in some areas of the *Nfat1*^−/−^ synovium (***Figure 2A-B**, Synovium*). Pathological chondrogenesis and endochondral ossification also occurred in the subchondral bone leading to localized thickening of the subchondral bone in the *Nfat1*^−/−^ facet joints, but not in the age-matched WT facet joints (***Figure 2A-B**, Sub-bone*). Cartilage formation with endochondral ossification in the joint margins was first observed at 6 months, which eventually formed chondro-osteophytes or osteophytes at 12 and 18 months in the *Nfat1*^−/−^ facet joints, but not in the age-matched WT facet joints (***Figure 2A-B**, Joint margin*).

**Figure 2.**
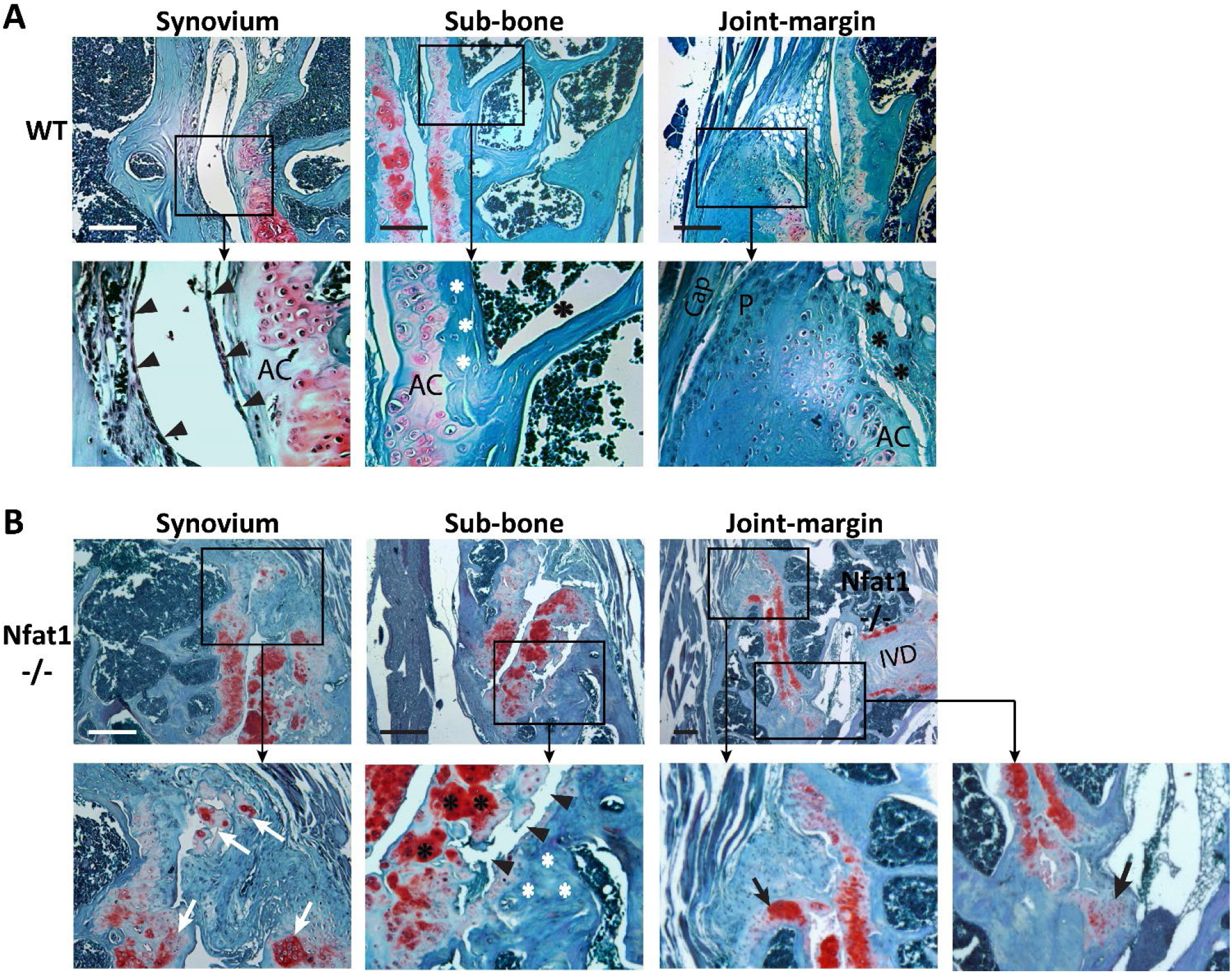
Osteoarthritic changes in the peri-articular tissues of NFAT1-mutant (Nfat1−/−) facet joints. (**A**) Facet joints from 18-month-old WT mice show essentially normal synovium with synovial cavity, synovial lining cells (black arrowheads), subchondral bone (Sub-bone) with cortical bone plate (white *), joint margin containing capsule (Cap), periosteum (P) and synovium (black *), as well as articular cartilage (AC) with mild degeneration. (**B**) Facet joints from 18-month-old Nfat1−/− mice show severe OA changes in the synovium with ectopic cartilage formation (white arrows), subchondral bone thickening (white *), cartilage-bone separation (black arrowheads), and joint margin with chondro-osteophyte formation (black arrows), IVD = intervertebral disc. Safranin-O and fast green staining, scale bar = 100 μm.

In general, FJOA occurred bilaterally and OA changes in the facet joints of lumbar vertebrae 4-5 (L4-5), L5-L6 and L6-S1 were more severe than L3-L4 in *Nfat1*^−/−^ mice. Six mice per genotype (*Nfat1^−/−^* vs WT), both male and female, were examined at each age/time point. Gender differences in histopathology were not observed.

### Development and reproducibility analysis of a novel FJOA scoring system

A novel semi-quantitative scoring system (***Table 1A–B***) was developed to objectively determine the age-dependent severity of FJOA in NFAT1-mutant and WT (control) mice. Coronal sections of the lumbar spine samples stained with safranin-O and fast green were used for imaging which covered articular cartilage, joint margin, subchondral bone, synovium, joint capsule, and surrounding muscles of the facet joints. Statistical analysis of inter-observer and intra-observer variability using Pearson’s correlation coefficient tests demonstrated low inter- and intra-observer variabilities, indicating the high reproducibility and ease of use of our novel FJOA scoring system by three observers/scorers at different levels of OA research experience (***Figure 3A-B***). Tissue specific scoring revealed that cartilage scores across the highly experienced, experienced, and novice observers had the greatest consistency with no variations in any measurements at any age points, while osteophyte and synovitis scores among the observers showed significant variation in only two measurements: osteophyte scores at 18 months of the first measurement (***Supplementary Figures* 1)** and synovitis scores at 12 months of the second measurement (***Supplementary Figures 2***).

**Figure 3.**
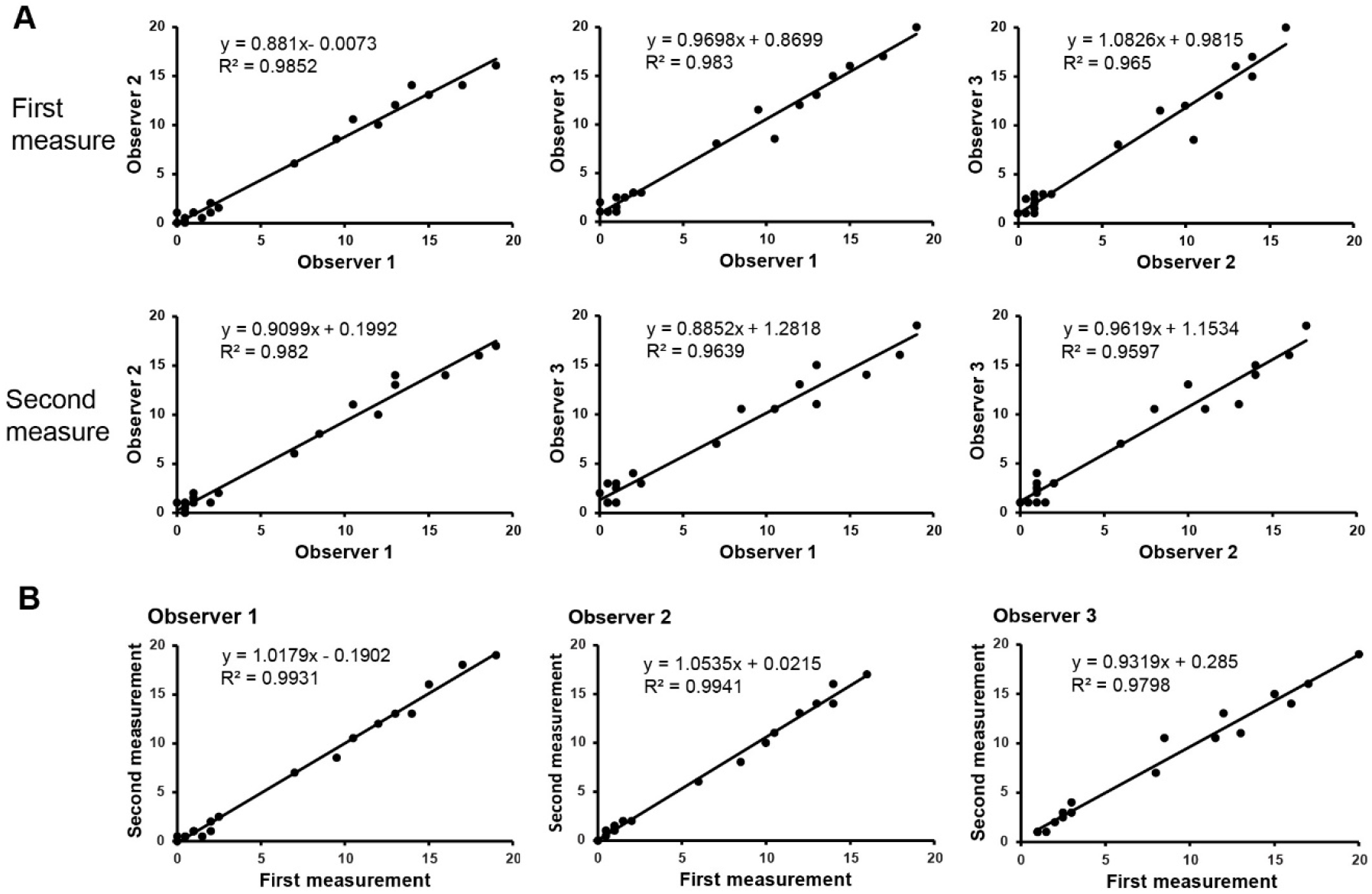
Inter- and intra-observer variabilities of the FJOA scoring system. (**A**) Inter-observer variability tests with linear regression analysis demonstrate low inter-observer variabilities among the three observers (scorers) and high reproducibility of the newly developed FJOA scoring system for both the first and second measurements (measure). (**B**) Intra-observer variability tests indicate low intra-observer variabilities between two measurements with a minimum interval of one week by the same individual.

**Table 1A.** Semi-quantitative FJOA scoring system: articular cartilage

| Score | Articular cartilage lesion (per facet*) |
| --- | --- |
| 0 | None |
| 0.5 | Focal loss of Safranin-O staining without structural changes |
| 1 | Surface abrasion, fissuring, or fibrillation within the superficial layer |
| 2 | Vertical clefts down to the layer immediately below the superficial layer with/without loss of surface lamina |
| 3 | Vertical clefts/erosion to the calcified cartilage extending to <50% of the articular surface |
| 4 | Vertical clefts/erosion to the calcified cartilage extending to >50% of the articular surface |
| 5 | <50% Cartilage loss in articular surface width with erosion to subchondral bone |
| 6 | >50% Cartilage loss in articular surface width with erosion to subchondral bone |

**Table 1B.** Semi-quantitative FJOA scoring system: periarticular tissues

| Score | Chondro-osteophyte (per<br>facet*) | Score | Subchondral bone<br>change (per facet*) | Score | Synovitis (per joint<br>end**) |
| --- | --- | --- | --- | --- | --- |
| 0 | None | 0 | None | 0 | None |
| 1 | Chondro-osteophyte present<br>either proximal or<br>distal joint margin | 1 | Subchondral bone<br>thickening ($\geq 2$-fold<br>of normal) | 1 | Thickening of synovium<br>($\geq 2$-fold of normal<br>synovial thickness) |
| 2 | Chondro-osteophyte present<br>both proximal and<br>distal joint margins | 2 | Subchondral bone<br>thickening with<br>reactive chondro-<br>genesis or cartilage-bone separation | 2 | Thickening of synovium<br>with ectopic<br>chondrogenesis and/or<br>ossification in<br>subintimal stroma |

\*Score from each facet (either inferior or superior) of the facet joint
\*\*Score from each joint end (either cephalic/proximal or distal portion of the facet joint)

### Tissue- and facet-specific involvement in initiation and progression of FJOA

#### Tissue-specific scoring

Semi-quantitative histomorphometry using our newly developed FJOA scoring system with statistical analyses demonstrated a slow but significant progression of overall OA severity in *Nfat1*^−/−^ facet joints from 2 to 18 months of age (***Figure 4A***). Tissue-specific scoring revealed significantly increased cartilage degradation in *Nfat1*^−/−^ facet joints as early as 2 months when compared with WT facet joints. In contrast, no significant differences in chondro-osteophyte (Osteophyte), subchondral bone (Sub-bone) changes, and synovitis were observed between *Nfat1*^−/−^ and WT facet joints at 2 months. Age-related progression analysis showed significantly increased cartilage lesions, osteophyte formation, and subchondral bone changes between 2 and 6 months, significantly increased cartilage lesions between 6 and 12 months, and significantly more severe osteophyte formation and synovitis between 12 and 18 months (***Figure 4B-E***).

**Figure 4.**
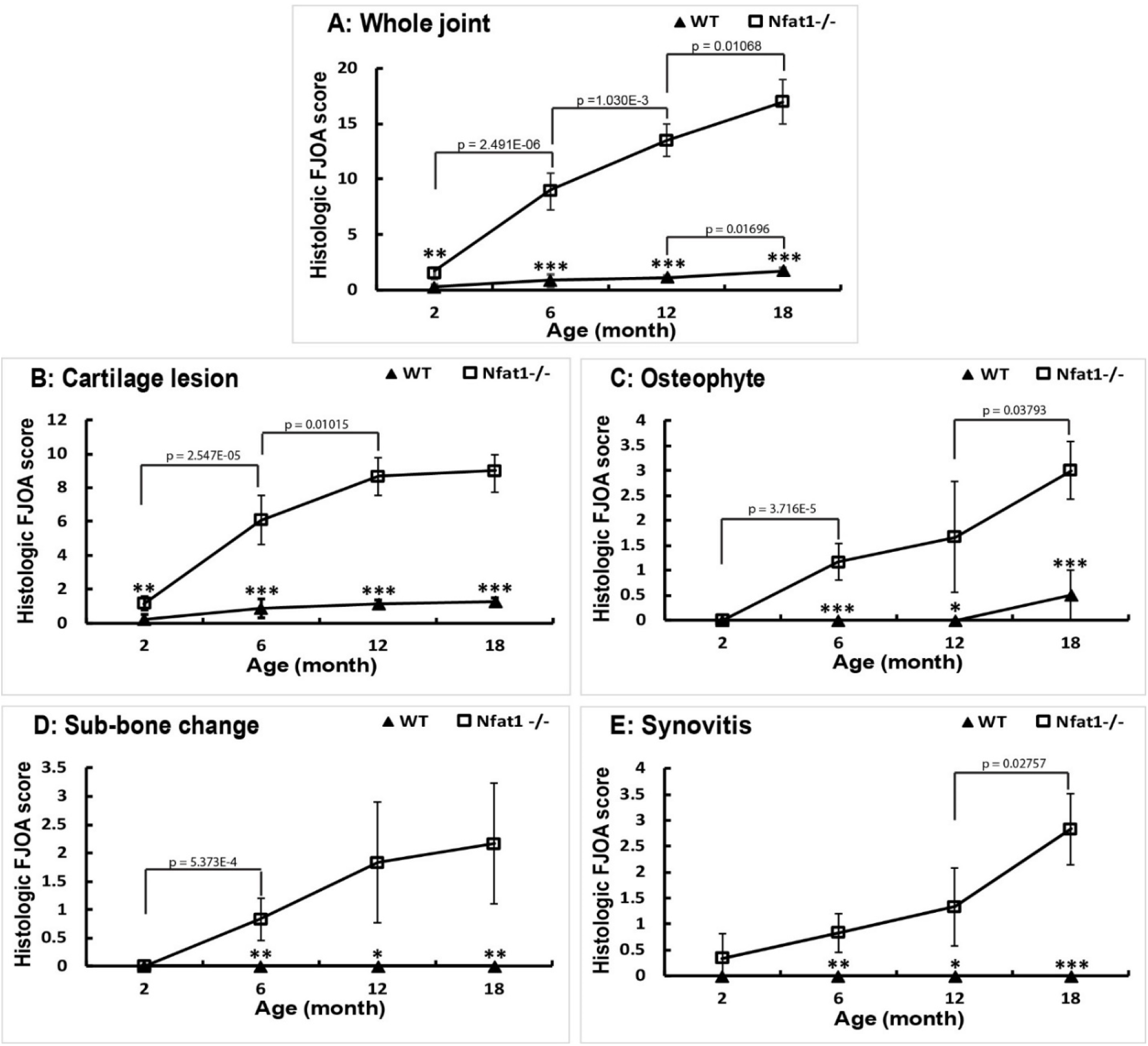
Age-dependent progression of FJOA severity for the whole joint and area-specific osteoarthritic changes. The histologic FJOA scores presented here were the mean ± SD of the examined joints. The levels of error-bar represent the deviations among the different images of the same genotype/age group, not the deviations among the observers. (**A**) The whole joint OA scores are presented using an averaged summed score at each age point. (**B-E**) The histologic FJOA score for specific osteoarthritic characteristics including cartilage lesion, chondro-osteophyte formation, subchondral bone (Sub-bone) change, and synovitis. For **A-E**, the p values represent the statistical significance between the age points of Nfat1−/− facet joints. The * represents the statistical difference between WT and Nfat1−/− facet joints at the same age point. *p < 0.05, ** p < 0.01, ***p < 0.001.

#### Facet-specific scoring

To determine the progression rate of FJOA in specific facets of the joint, we conducted facet-specific histomorphometry analysis using the new FJOA scoring system with statistical analyses. From 2 to 6 months, cartilage lesions were significantly progressed in both the inferior (inf) and the superior (sup) facets, osteophyte formation was significantly progressed in the inferior facet but not the superior facet, subchondral bone changes significantly progressed in the superior facet but not the inferior facet, and synovitis significantly progressed in the distal (dist) end but not in the proximal (prox) end. From 6-12 months, cartilage lesions were significantly progressed in the inferior facet but not the superior facet. From 12 to 18 months, osteophyte formation significantly progressed in the superior facet but not the inferior facet, while synovitis became more severe in the distal end but not in the proximal end (***Supplementary Figure 3***).

### Aberrant gene/protein expression in both cartilage and synovium of *Nfat1*^−/−^ facet joints at the initiation stage of FJOA

At 2 months of age, qPCR analysis detected significantly decreased expression of *Acan* mRNA (encoding aggrecan) with significantly increased expression of *Il1b* mRNA (encoding interleukin-1β/IL-1β) in the *Nfat1*^−/−^ facet joint cartilage when compared with WT facet cartilage (***Figure 5A***). Significantly increased expression of *Mmp3* (encoding matrix metalloproteinase-3/MMP-3), *Il1b* and *Tnfa* (encoding tumor necrosis factor-α/TNF-α) was detected in the *Nfat1*^−/−^ facet joint synovium (***Figure 5B***). Immunohistochemistry (IHC) demonstrated overexpression of the IL-1β and MMP-3 proteins in *Nfat1*^−/−^ facet cartilage and synovium, while the TNF-α protein was primarily overexpressed in *Nfat1*^−/−^ facet synovium (***Figure 6***). These findings suggest that dysfunction of chondrocytes and synovial cells contributes to the initiation of spontaneous FJOA and that molecular biological alterations occur much earlier than structural OA changes in this FJOA model, which provide new insights into the molecular and cellular mechanisms of spontaneous FJOA.

**Figure 5.**
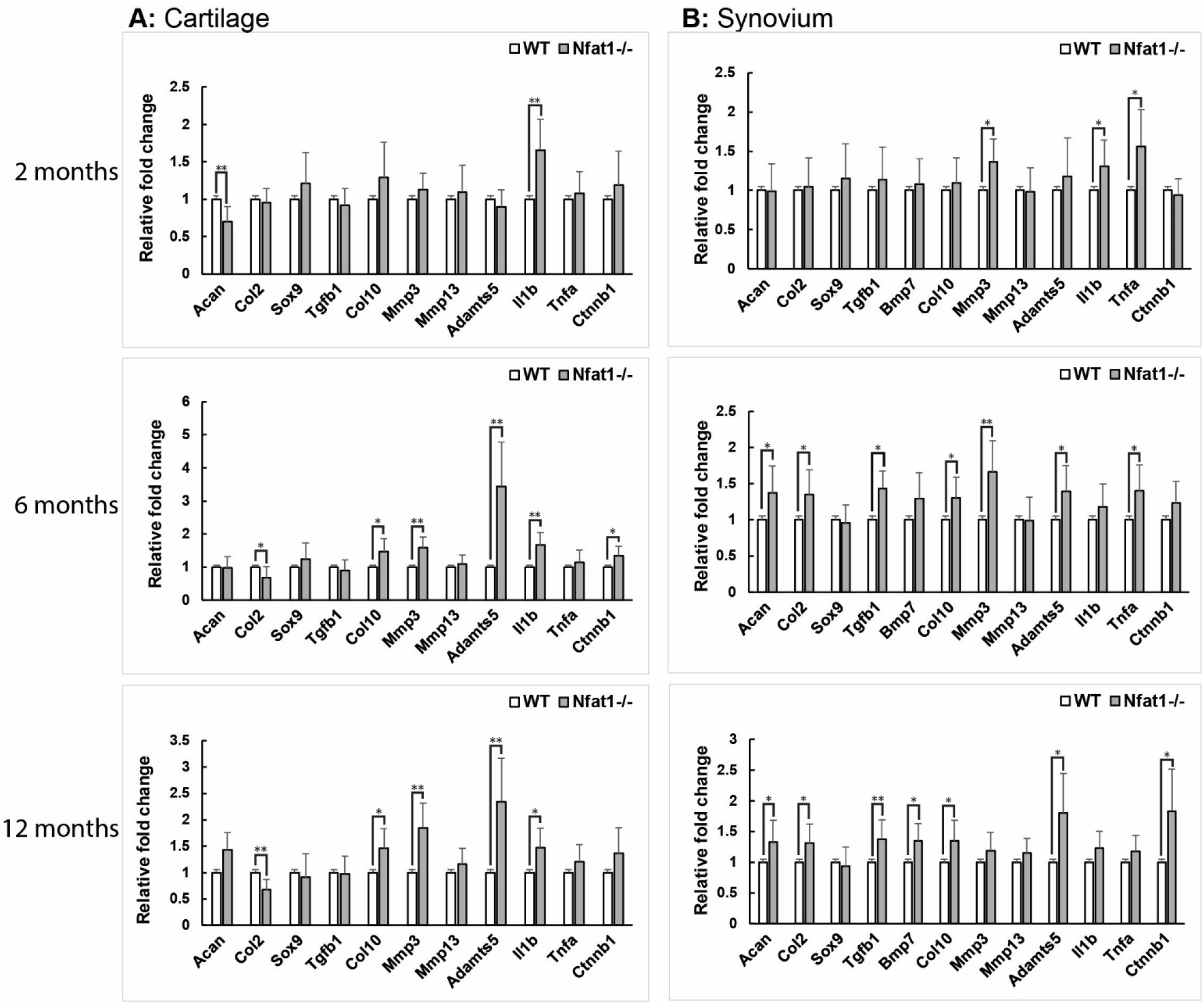
qPCR analysis showing differential expression of anabolic and catabolic genes in facet joint cartilage and synovium from WT and Nfat1−/− mice at 2, 6, and 12 months of age. The expression level of each wild-type group has been normalized to “1.0”. N = 6. *p < 0.05, **p < 0 .01.

**Figure 6.**
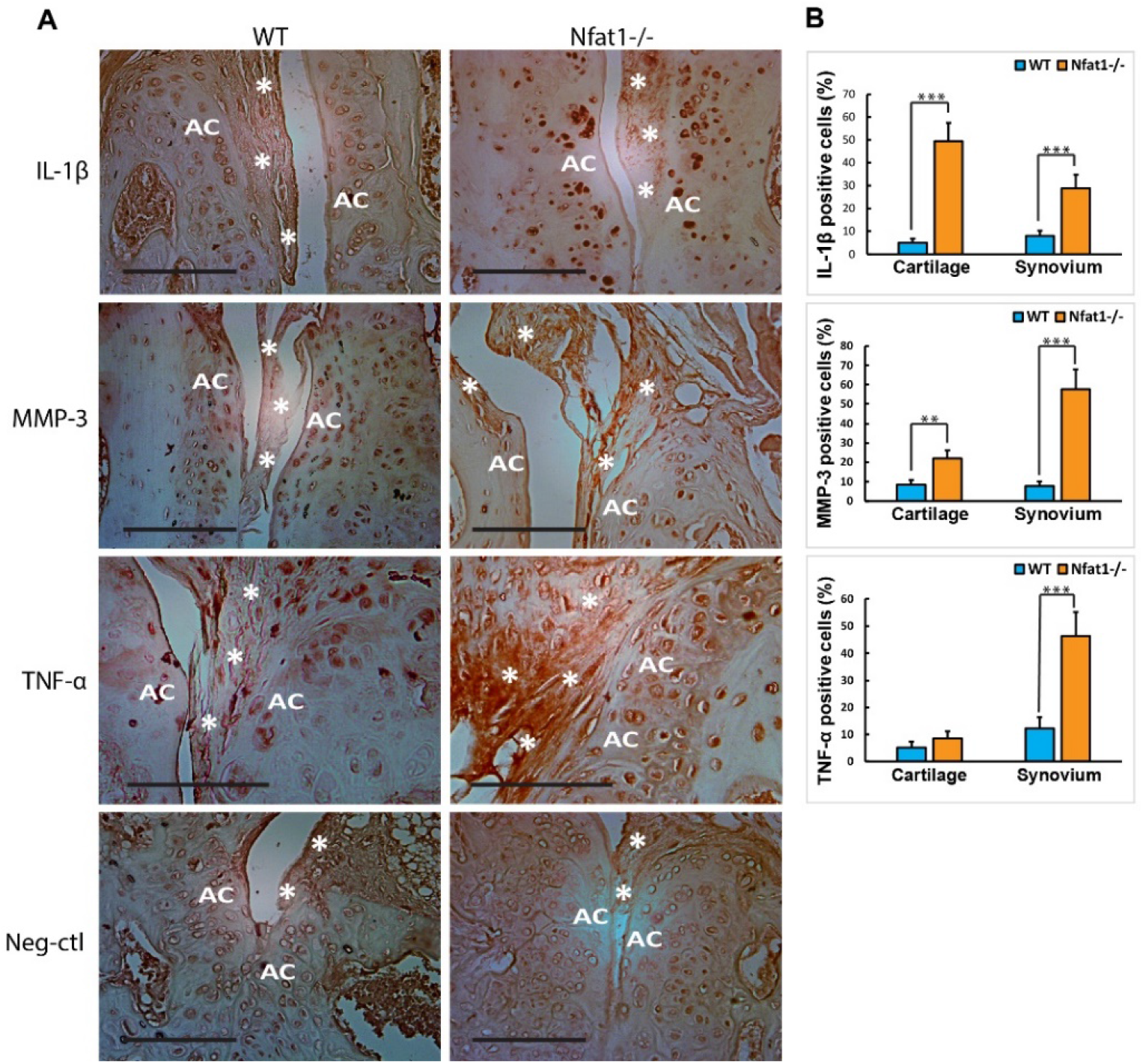
Representative immunohistochemistry (IHC) images of WT and NFAT1-mutnt (Nfat1−/−) facet joints at 2 months of age. (**A**) IHC images show increased expression of IL-1β, MMP-3 and THF-α in the cartilage and/or synovium of Nfat1−/− facet joints. Neg-ctl = negative control without using primary antibodies, AC = articular cartilage, white*denotes synovium. IHC positive cells and matrices are stained in brown with DAB chromogen. IHC with hematoxylin counterstaining, scale bar = 100 μm. Primary antibodies used for IHC: monoclonal IL-1β antibody (Santa Cruz, SC-52012), monoclonal MMP-3 antibody (LSBio, LS-C800335), and monoclonal TNF-α antibody (Santa Cruz, SC-52746). (**B**) Quantification of the percentage of cells positive for IL-1β, MMP-3 and THF-α, respectively, from IHC images (n = 4 per group). Significant differences in the number of positive cells between the WT and Nfat−/− groups were determined using two-tailed Student’s t-tests. **P<0.01; ***p<0.001.

### Temporal and spatial differences in gene expression changes during the progression of FJOA

qPCR analysis revealed aberrant gene expression in the cartilage of *Nfat1*^−/−^ facet joints in an OA-stage dependent manner (temporal differences). At 2 months of age, significantly decreased expression of *Acan* with significantly increased expression of *Il1b* was detected in the *Nfat1*^−/−^ facet cartilage when compared with WT facet cartilage. At 6 months, significantly decreased expression of *Col2a1* (encoding collagen-2) and significantly increased expression of *Col10a1* (encoding collagen-10), *Mmp3*, *Adamts5* (encoding a disintegrin and metalloproteinase with thrombospondin motifs-5), *Il1b*, and *Ctnnb1* (encoding β-catenin) were detected in the *Nfat1*^−/−^ facet cartilage. At 12 months, significantly decreased expression of *Col2a1* and significantly increased expression of *Col10a1*, *Mmp3*, *Adamts5*, and *Il1b* were detected in the *Nfat1*^−/−^ facet cartilage (***Figure 5A***).

Temporal differences in gene expression was also detected in the synovial tissue of *Nfat1*^−/−^ facet joints. qPCR analysis revealed significantly increased *Mmp3, Il1b* and *Tnfa* at 2 months, significantly increased *Acan, Col2a1, Tgfb1, Col10a1* (a marker of hypertrophic chondrocytes), *Mmp3*, *Adamts5* and *Tnfa* at 6 months, and significantly increased *Acan*, *Col2a1*, *Tgfb1*, *Bmp7* (encoding bone morphogenetic protein-7), *Col10a1*, *Adamts5* and *Ctnnb1* at 12 months (***Figure 5B***). Members of the TGF-β/BMP superfamily may induce chondrogenic differentiation of mesenchymal stem cells in the synovium.^28^ An increase in *Acan*, *Col2a1* and *Col10a1* and *Ctnnb1* expression in the *Nfat1*^−/−^ synovium reflects ectopic chondrogenesis and increased chondrocyte hypertrophy with pathologic endochondral ossification. The temporal differences in expression of a specific gene suggest that molecular changes in the joint tissue cells are a dynamic process during the progression of FJOA.

Spatial differences in expression levels of specific genes between cartilage and synovium were remarkable. For example, *Col2a1* expression was significantly decreased in *Nfat1*^−/−^ cartilage at 6 and 12 months due to cartilage degradation but was significantly increased in *Nfat1*^−/−^ synovium at 6 and 12 months reflecting pathological chondrogenesis in the *Nfat1*^−/−^ facet synovium. *Tnfa* expression was significantly upregulated in synovium, but not in cartilage. The tissue-specific gene expression patterns suggest that different types of cells in these tissues may play distinct roles during the initiation and progression of FJOA.

The evidence-based molecular and cellular mechanisms for initiation and progression of the NFAT1 mutation-mediated spontaneous FJOA, as well as possible involvement of mechanical stress secondary to OA structural changes, are summarized in ***Figure 7***.

**Figure 7.**
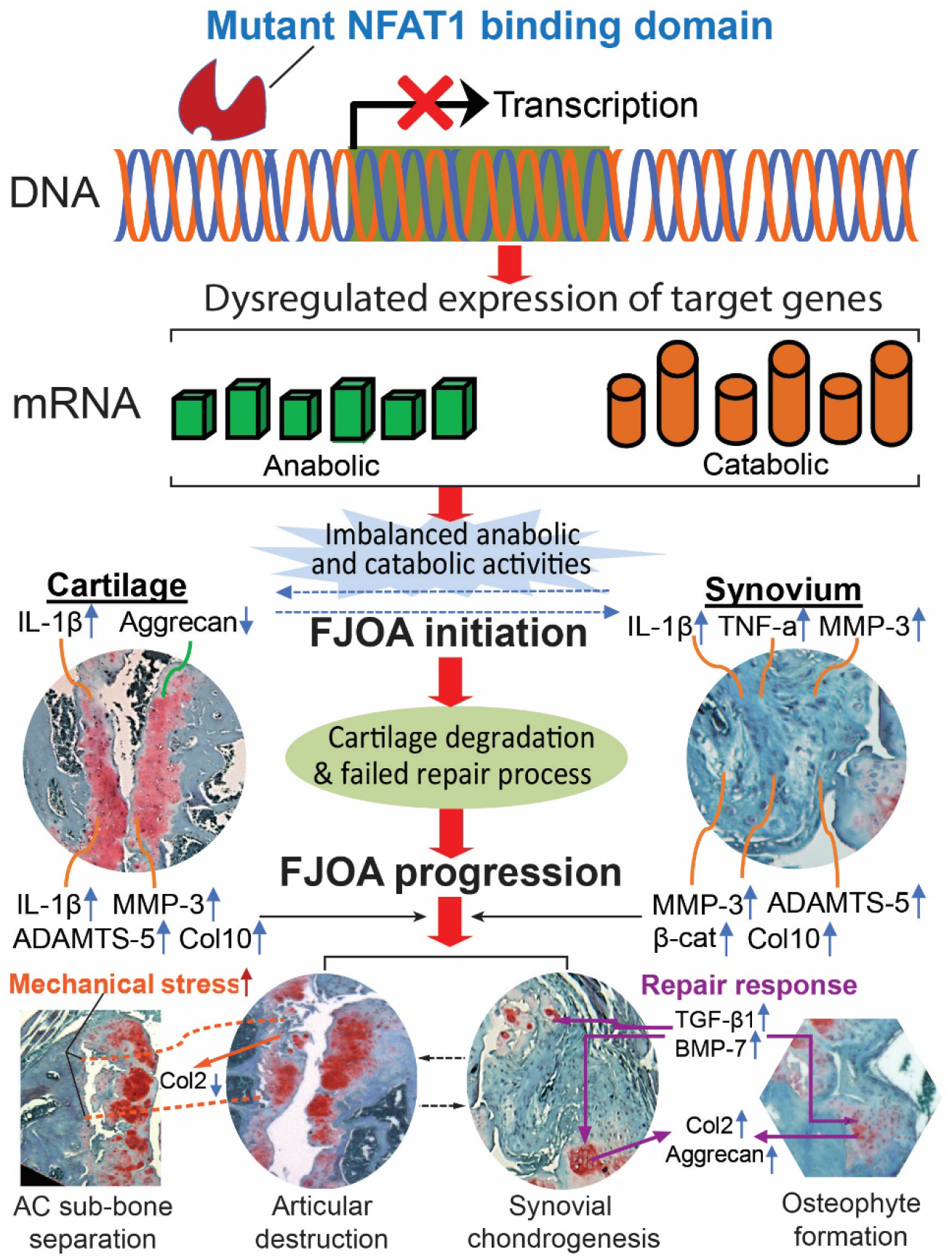
A schematic summary of pathogenic mechanisms for the NFAT1 mutation-mediated spontaneous FJOA. Arrows with solid lines indicate mechanistic observations from authors’ studies. Arrows with dotted lines indicate possible FJOA and general OA mechanisms proposed by others ^4,34^. The mutant NFAT1 binding domain compromises its binding to the promoter of target genes, resulting in dysregulated expression of NFAT1 target genes in joint tissue cells and imbalanced anabolic and catabolic activities in facet joint cartilage and synovium. These changes lead to cartilage degradation and trigger reparative reactions in joint tissues, particularly in the synovium, resulting in upregulated expression of anabolic factors (e.g. TGF-β1 and BMP-7) and causing ectopic chondrogenesis and chondro-osteophyte formation. However, the reparative response fails to halt the progression of FJOA due to continued presence of pathogenic factors as well as increased mechanical stress on the cartilage and subchondral bone. AC = articular cartilage, sub-bone = subchondral bone.

## DISCUSSION

The current study has characterized the first murine model of spontaneous FJOA with age-dependent osteoarthritic characteristics in multiple joint tissues. To our knowledge, it is also the first animal model of spontaneous FJOA. Animal models of OA can be classified as mechanically or surgically induced, chemically induced, and spontaneous (naturally occurring) OA. Surgically or chemically induced OA animal models requires invasive local manipulations to produce mostly unilateral OA with acute joint inflammation and rapid disease progression. ^15–19^ In contrast, spontaneous OA animal models develop without invasive local manipulations but may be mediated by systemic genetic predisposition, which usually occur bilaterally with slow progression, resembling primary or idiopathic OA in humans. ^29–31^ The STR/ort mouse with a piebald mutation is a widely used spontaneous mouse OA model developed by carcinogen treatment; additional spontaneous animal OA models were developed using transgenic technologies.^29,31–33^ To our knowledge, those previously reported spontaneous OA models were exclusively focused on the appendicular synovial joints, but not on the facet joints. Therefore, the NFAT1-mutant mouse represents the first characterized animal model of spontaneous FJOA.

Also, for the first time, the present study has successfully developed a histopathologic FJOA scoring system to semi-quantitatively measure the tissue-specific severity of FJOA and age-related progression of the disease. The development of the novel scoring system mainly attribute to the various degrees of histopathologic features seen in the *Nfat1*^−/−^ facet joints, from mild articular surface abrasion to severe OA changes such as deep cartilage clefts, loss of cartilage, thickening of subchondral bone, separation of degraded cartilage from the subchondral bone, remarkable chondro-osteophytes, and synovitis with ectopic chondrogenesis in the synovium and capsule. Our careful literature review suggests that such severe FJOA does not appear to be present in the previously reported animal models of FJOA induced by invasive local manipulations in which only mild to moderate cartilage degeneration was observed, with little or no articular destruction or osteoarthritic changes in the periarticular tissues ^17,18,20^. OA, including FJOA, is the most common form of joint disease. Although many risk factors and therapeutic approaches have been tested, no disease-modifying pharmacologic therapy is currently available to stop or reverse the progression of OA largely because the pathogenic mechanisms of OA remain unclear.^34–36^ The present study has provided new insights into the pathogenesis of FJOA. First, it has been a long debate whether OA pathogenesis starts with cartilage degradation, subchondral bone alterations, or osteochondral junction abnormalities. ^34,37,38^ The histological and molecular findings from this study suggest that dysfunction of articular chondrocytes and synovial cells contributes to the initiation of spontaneous FJOA, and that changes in the subchondral bone and osteochondral junction observed in the later stages of FJOA are likely to be secondary alterations. Second, the current study demonstrated the temporal and spatial differences in expression levels of specific genes in facet joint cartilage and synovium as shown in Figure 5. These findings suggest that FJOA progression is a dynamic process in specific joint tissues. Thus, an expression level of a single matrix protein, cytokine, or proteinase in a single join tissue at a single time point should not be used as a specific biomarker for diagnosis or assessment of therapeutic efficacy of FJOA.

The present study has also provided new insights into the therapeutic strategies for FJOA and general OA. First, although OA has been recognized as a disease of the joint as an organ^39^, previous and ongoing OA clinical trials mainly target articular cartilage with little attention to synovium. This study demonstrated that synovial cells in the NFAT1-mutant facet joints release cartilage-degrading catabolic mediators as early as 2 months of age, well before the osteoarthritic structural changes, indicating that dysfunction of synovial cells is involved in the early initiation of FJOA. Since inflamed synovium is responsible for cartilage breakdown and is an important source of joint pain ^34^, OA therapies targeting both synovium and cartilage would more effectively attenuate cartilage degradation, joint pain, and overall OA progression. Second, many OA clinical trials targeting a single catabolic molecule have been unsuccessful due to poor efficacy. ^40–42^ The present study demonstrated abnormal expression of multiple anabolic (e.g. *Col2a1*, *Acan, Tgfb1, and Bmp7*) and catabolic (e.g. *Il-1b*, *Tnfa*, *Mmp3*, *Adamts5*, and *Col10a1*) genes in both cartilage and synovium of *Nfat1*^−/−^ facet joints. All these genes are potential NFAT1 target genes whose expression is transcriptionally regulated by the transcription factor NFAT1. This is confirmed by our previous study showing that NFAT1 can directly bind to at least 21 genes encoding cartilage-matrix proteins, growth factors, inflammatory cytokines, matrix-degrading proteinases, and specific transcriptional activators.^24^ Since NFAT1 is an upstream factor regulating the expression of multiple anabolic and catabolic genes in both cartilage and synovium, NFAT1 could be a more promising molecular target for OA therapy than other anti-OA drug candidates that target a single, downstream anabolic or catabolic molecule.

FJOA is a major source of low back pain because the facet joints are highly innervated with rich nociceptive fibers. FJOA, along with IDD, is a leading cause of chronic disability in humans. Management of FJOA remains a difficult challenge for the physician caring for patients with spinal disorders.^7,8,10,43^ Low back pain alone causes more global disability than any other health condition.^44^ The spontaneous FJOA animal model identified in the present study has substantially advanced our understanding of the mechanisms and characteristics of FJOA and could help us develop novel preventive and therapeutic strategies for FJOA and low back pain.

A limitation of this study is that experimental evaluations are focused on the histopathologic and molecular biologic analyses without pain assessments, mainly due to scientific concerns. Vocalization threshold testing, one of the pain assessments for low back pain with algometry on the skin over the facet joint region ^17^, was not conducted in this study. This is because vocalization testing is mainly used for measuring the difference between the paired facet joints, which works for unilateral disorder but is not suitable for bilateral facet disorders. The current study does not include alterations in the intervertebral disc and other spinal structures of NFAT1-mutant mice, which will be analyzed and reported separately.

In conclusion, the present study has identified and carefully characterized the first murine model of spontaneous FJOA and developed the first FJOA scoring system. Our molecular and histomorphometric analyses have revealed that the NFAT1 mutation in the DNA binding domain causes dysregulated transcription of multiple anabolic and catabolic genes in the facet cartilage and synovium, thereby initiating a slow-progressing FJOA. This suggests that NFAT1 is essential for maintaining the homeostasis of cartilage and synovium of the facet joints. These novel findings have provided new insights into the pathogenetic mechanisms of FJOA, both in animal models and in humans. The results of this pre-clinical study warrant further investigation to explore if a disruptive NFAT1 mutation or a significant decrease in NFAT1 expression level is a risk factor for development of FJOA in humans.

## MATERIALS AND METHODS

### Animals

Both genders of 2-18 months old BALB/c background mice bearing mutant NFAT1 alleles (NFAT1-mutant mice) and their wild-type (WT) littermates were used for this study. The mutant mice were generated by deleting an exon of the *Nfat1* gene encoding amino acids of the DNA binding domain. The methods for generation of the initial NFAT1(NF-ATp)-mutant mice were described previously.^22,23^ The original breeders of NFAT1-mutant mice used in this study were a gift from Dr. Laurie Glimcher (Harvard University). Mice homozygous for the disrupted allele are referred to as NFAT1-mutant or *Nfat1^−/−^* mice in this study. The genotype of *Nfat1^−/−^* mice was confirmed by genotyping using specific primers. All animal procedures were approved by the Institutional Animal Care and Use Committee in compliance with federal and state laws and regulations.

### Histology, histochemistry, and immunohistochemistry (IHC)

After euthanasia of the experimental animals, spine tissue samples covering the segments of lumbar 3 to sacral 1 (L3-S1) vertebrae and facet joints were harvested from *Nfat1^−/−^* and WT mice (negative controls) at the ages of 2, 6, 12, and 18 months for histopathological analyses. We chose to examine L3-S1 lumbar tissue samples because FJOA most commonly occurs in these spinal levels in humans. Harvested lumbar spine samples were fixed in 2% paraformaldehyde, decalcified in 25% formic acid, embedded in paraffin, and sectioned at 5 μm. The tissue sections were stained with safranin-O and fast green to identify cartilage cells and matrices, as well as haematoxylin and eosin (H&E) for structural and cellular analyses of bone and soft tissues. Tissue sections that had been decalcified in 10% EDTA were used for IHC using specific antibodies to identify the location and intensity of the proteins of interest. To observe both immunoreaction and cellular morphology, some tissue sections were stained by the avidin-biotin peroxidase complex methods. DAB chromogen was used for color detection as described previously. ^24,25,45^ Six animals per genotype/gender/age group were utilized for histological analysis.

### Development of a novel FJOA scoring system

A novel semi-quantitative scoring system was developed to determine the severity of FJOA. Coronal sections of the lumbar spine samples stained with safranin-O and fast green were used for imaging which covered articular cartilage, joint margin, subchondral bone, synovium, capsule, and surrounding muscles of the facet joints. A set of 20 images of both WT and *Nfat1^−/−^* lumbar facet joints were scored twice with a minimum time interval of 1 week to obtain intra-observer variability from the same observer. The images were scored by a highly experienced observer (JW, Observer 1), an experienced observer (MM, Observer 2), and a novice observer (XL, Observer 3) with no previous experience in OA histology to assess inter-observer variability as well as to determine the ease of use of the scoring system.

### Histomorphometry

To determine FJOA severity, the facet joint at the L5/L6 level (the mouse has 6 lumbar vertebrae ^46^) from each WT mouse and each *Nfat1^−/−^* mouse were utilized for histologic scoring because L5/L6 *Nfat1^−/−^* facet joints usually displayed the most severe OA among the examined lumbar segments. OA histologic scoring was conducted by 2 independent observers who were blinded regarding genotypes and ages. The maximum OA score for each fact joint was set at 24, which covered cartilage lesions (score range: 0-6 per facet, 0-12 per joint), chondro-osteophyte formation (score range: 0-2 per facet, 0-4 per joint), subchondral bone changes (score range: 0-2 per facet, 0-4 per joint), and synovitis (score range: 0-2 per proximal (cephalic) or distal end, 0-4 per joint). A detailed semi-quantitative FJOA scoring system is presented in ***Table 1A–B***. Cumulative scores of a representative facet joint of each animal from both observers at each time point were averaged for statistical analysis to determine the genotype-specific and age-dependent changes in OA severity.

To determine the percentage of IHC positive cells in WT and *Nfat1^−/−^* facet joint cartilage and synovium, IHC images were analyzed with Fiji software system, an image processing package composed of ImageJ ^47^.

### Quantitative real-time reverse transcription polymerase chain reaction (qPCR)

Facet joint cartilage and synovium (containing a thin layer of fibrous capsular tissue due to technical difficulties in obtaining pure synovium from mouse facet joints) were freshly dissected under a microscope (LEICA M80, with a LEICA IC80 HD video camera). Cartilage and synovium samples were collected separately in RNAlater solution (Ambion, Austin, TX, USA) at the age of 2, 6, and 12 months. No tissue samples were collected for qPCR analysis at 18 months due to severe OA changes with articular destruction or fusion. Cartilage or synovium samples from L3-S1 facet joints of the same animal were pooled for RNA extraction in order to obtain enough RNA from each individual mouse. Tissue samples were homogenized in TRIzol reagent (Invitrogen, Carlslad, CA, USA) and treated with a DNA Digestion Kit (Ambion) to remove DNA. One microgram of total RNA and a High-Capacity cDNA Reverse Transcription Kit (Applied Biosystems, Foster City, CA, USA) were used to yield cDNA. Specific primers used in this study are presented in ***Supplementary Table 1***. qPCR reactions were performed in triplicate using a 7500 Real-Time qPCR system and SYBR Green reagents (Applied Biosystems). *Gapdh* expression levels were used as internal controls for load normalization of cDNA samples. Gene expression levels were quantified using 2^−ΔΔCt^ methods using the expression level of age-matched WT samples as controls as described previously.^24,25,48^

## Statistics

Quantitative data were presented as the mean ± standard deviation (SD). The significance of difference between means from two groups was analyzed by Student’s t-test; the difference between means for three or more groups was assessed by one-way ANOVA, followed by a post-hoc test (Tukey) using Excel 2017 software. Pearson’s correlation coefficient analysis was conducted using R: A Language and Environment for Statistical Computing software, Version 4.0.3, R Foundation for Statistical Computing. A p-value of less than 0.05 was considered statistically significant.

## Acknowledgments

This work was supported by the National Institute of Arthritis and Musculoskeletal and Skin Diseases of the National Institutes of Health (NIH) under Award Number R01 AR059088 (to JW) and the Mary & Paul Harrington Distinguished Professorship Endowment. Marc A. Asher, M.D., who made significant contributions to the design and performance of this study, has passed away after his lifelong devotion to orthopedic (spine) surgery and research.

## Author contributions

J.W. and M.A.A. conceived and designed the overall project; J.W., Q.L., M.J.M., and X.L. performed histopathologic and histomorphometric analyses; Q.L. conducted mouse genotyping; Y.F. performed microscopic tissue collection and qPCR gene expression analysis; X.L. conducted statistical analysis and figure preparation; D.C.B. and M.A.A. participated in data analysis and provided clinical input to the project; J.W. wrote the manuscript; Q.L., M.J.M., X.L., Y.F., D.C.B., and M.A.A. reviewed the manuscript.

---

**Supplementary Table 1.**
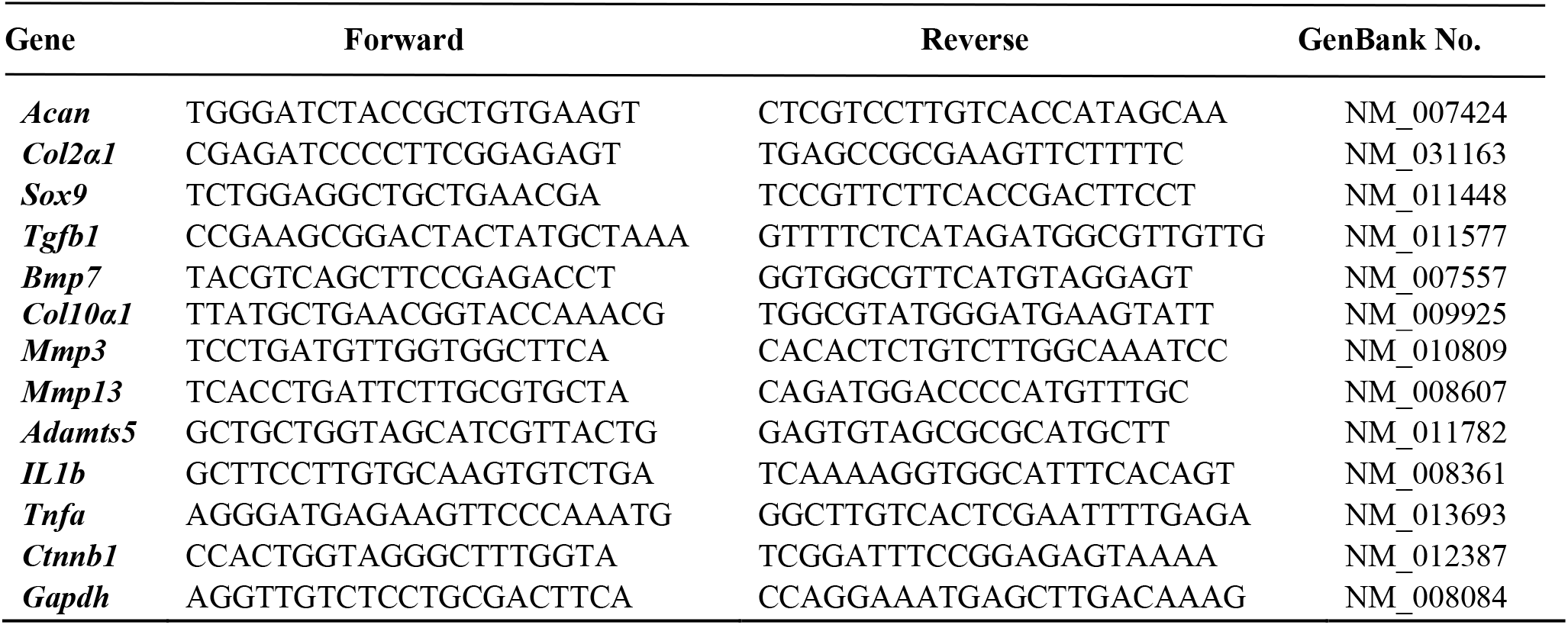
Specific primers used for qPCR analysis

**Supplementary Figure 1.**
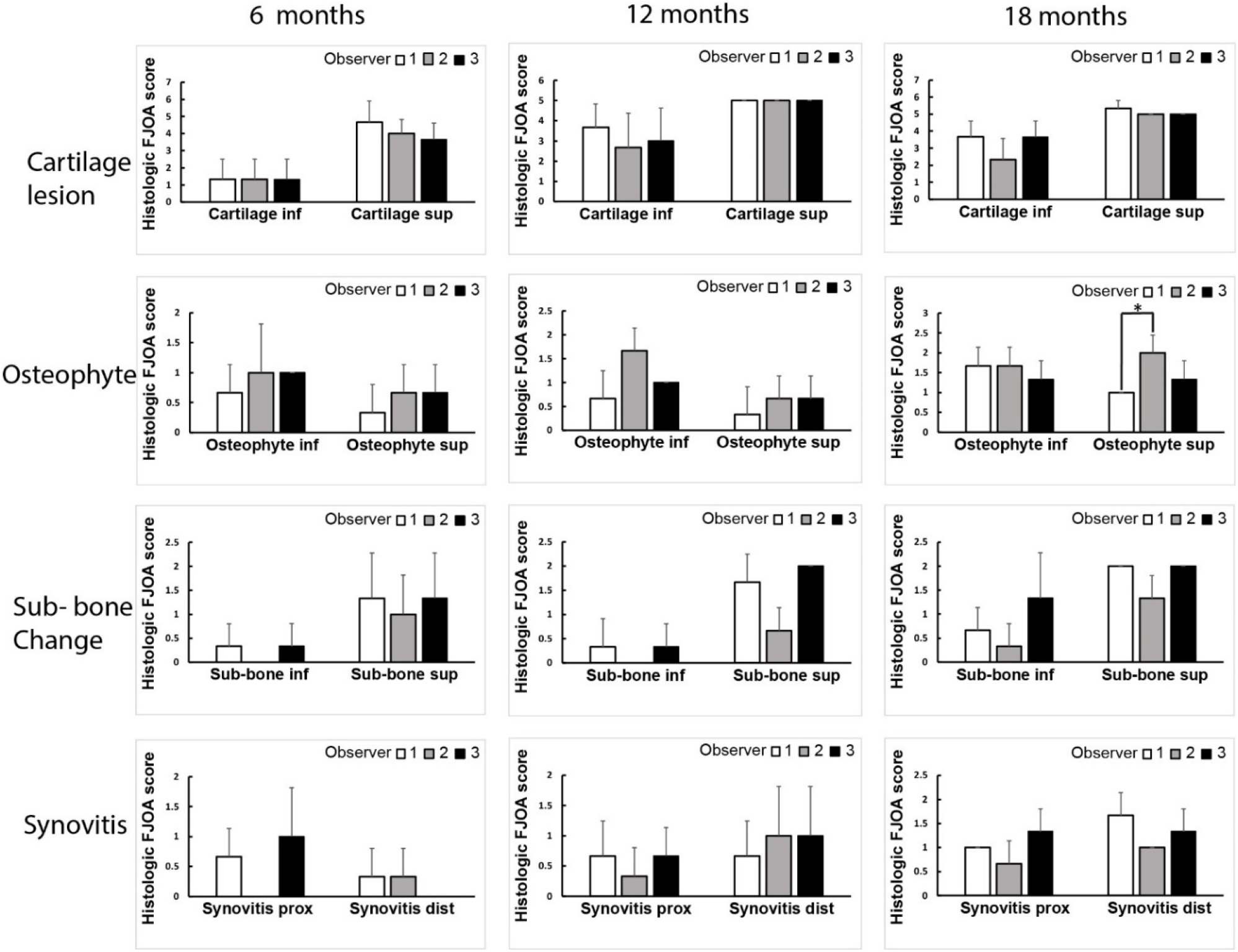
Comparative analyses of first measurement of area-specific inter-observer variabilities from three observers (Observer 1, 2, and 3) using the novel FJOA scoring system. Area-specific histologic FJOA scores for 2-month-old Nfat1−/− mice are not included as they are either none or too low to show. The histologic FJOA scores for specific areas from the three observers are highly reproducible with no significant differences among the observers, except the severity of osteophyte formation in the superior facet at 18 months between Observers 1 and 2. inf = inferior facet/articular process, sup = superior facet/articular process, Sub-bone = subchondral bone, prox = proximal/cephalic end of the facet joint, dis = distal/caudal end of the facet joint. *p < 0.05.

**Supplementary Figure 2.**
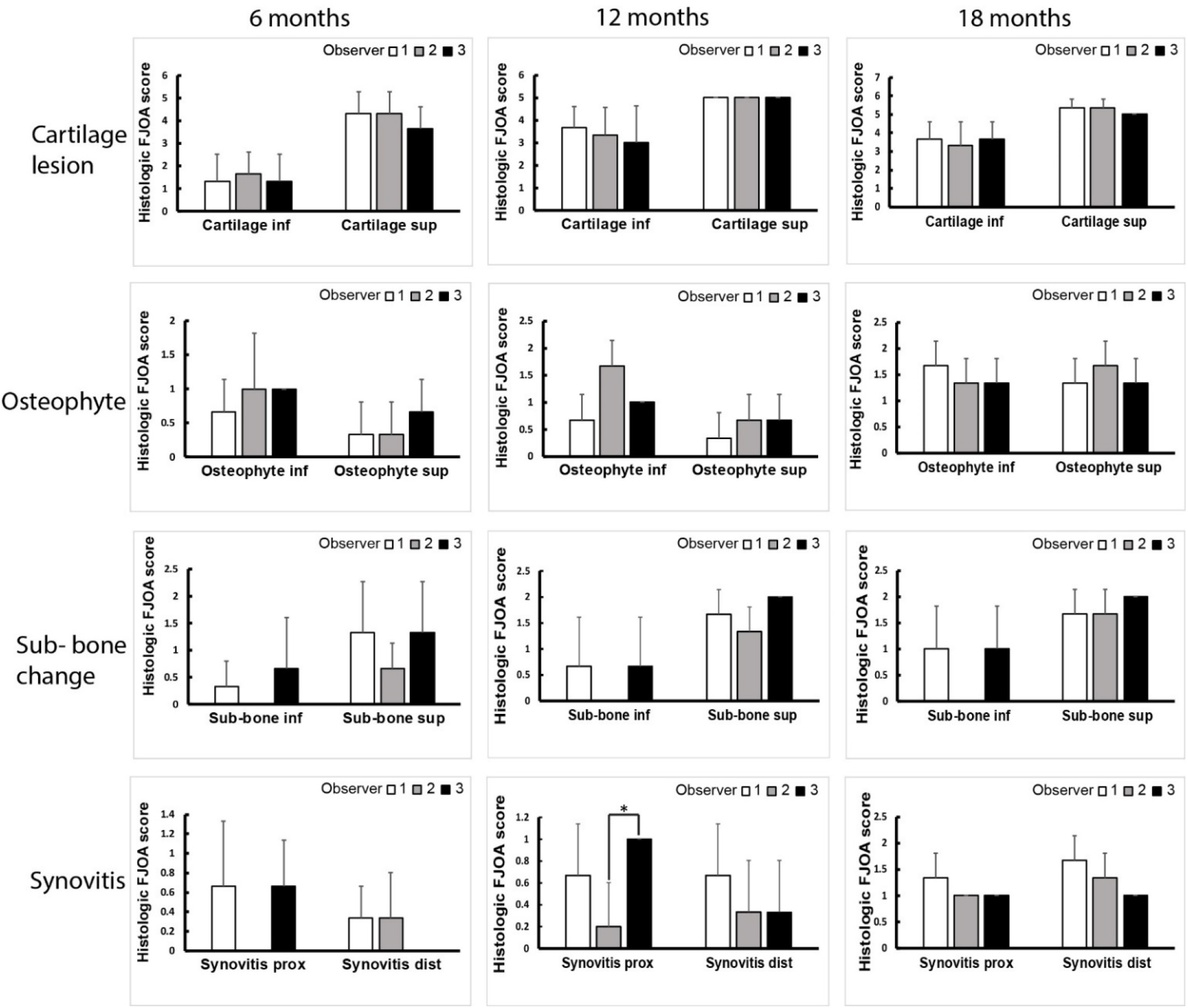
Comparative analyses of the second measurement of area-specific inter-observer variabilities from three observers (Observer 1, 2, and 3). Area-specific histologic FJOA scores for 2-month-old Nfat1−/− mice are not included as they are either none or too low to show. The histologic FJOA scores for specific areas from the three observers are highly reproducible with no significant differences among the observers, except the severity of synovitis in the proximal end of facet joints at 12 months, which showed a significant difference between Observers 2 and 3. inf = inferior facet/articular process, sup = superior facet/articular process, Sub-bone = subchondral bone, prox = proximal/cephalic end of the facet joint, dis = distal/caudal end of the facet joint. *p < 0.05.

**Supplementary Figure 3.**
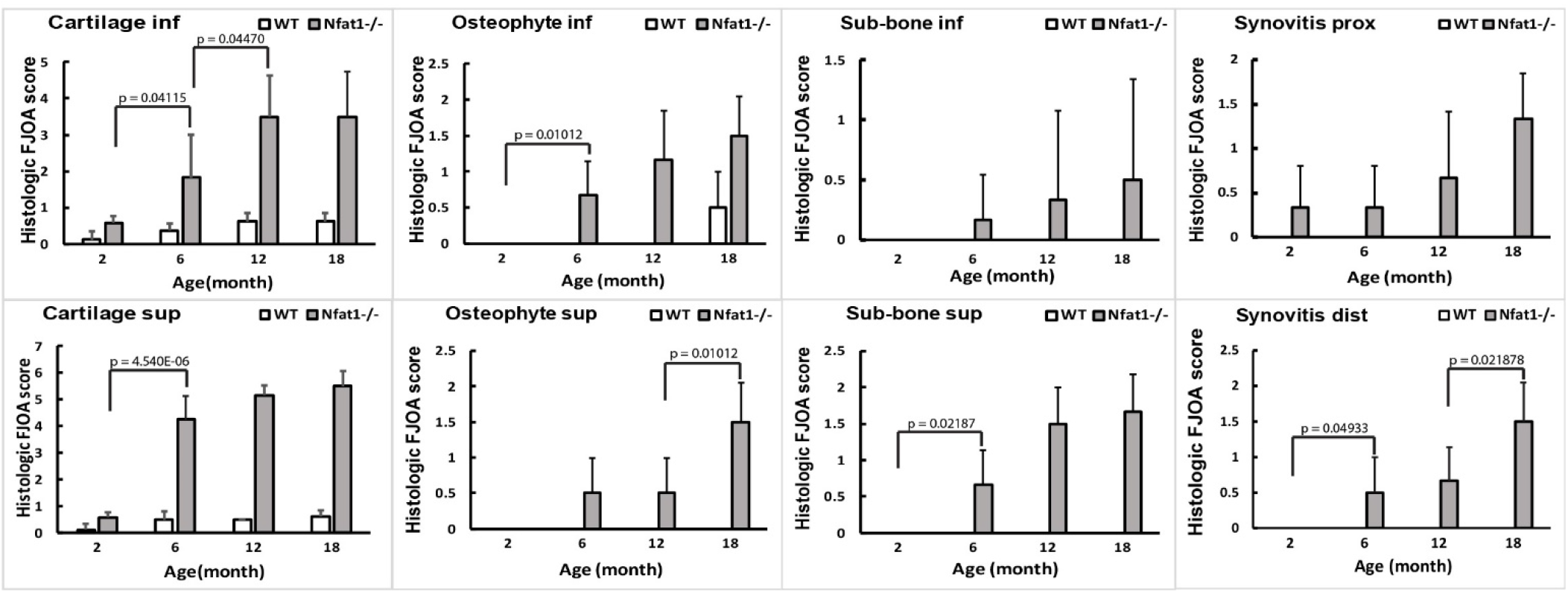
Age-related progression rates of FJOA in specific areas of the joint determined by histomorphometric analysis of FJOA severity. inf = inferior facet/articular process, sup = superior facet/articular process, Sub-bone = subchondral bone, prox = proximal/cephalic end of the facet joint, dis = distal/caudal end of the facet joint. The p values with statistical significance (p < 0.05) between two age points are indicated within the graphs.

## Notes

**Disclosure:** The authors declare no competing conflicts of interest.

### Competing Interest Statement

The authors have declared no competing interest.

